# Cyanophage thioredoxin and virulence enhancer genes differentially contribute to phage fitness and infection of marine *Synechococcus*

**DOI:** 10.1101/2025.10.05.680603

**Authors:** Ran Tahan, Dror Shitrit, Debbie Lindell

## Abstract

Environmental viruses, including marine cyanophages, carry many homologs of cellular genes (auxiliary metabolic genes), as well as small genes of unknown function (viral dark matter). However, the functional relevance and evolutionary benefits of these genes remain largely unknown. One such cellular-like gene codes for thioredoxin, a redox-regulating protein. Here we investigated the distribution of thioredoxin and a small gene of unknown function (*cve*) in diverse phage isolates. We found that thioredoxin is common among viruses isolated on diverse hosts from a variety of environments. In contrast, *cve* is restricted to marine cyanophages yet is present in multiple discrete cyanophage families. Investigation of the function of the thioredoxin gene via its deletion from a T7-like marine cyanophage, Syn5, revealed that it enhanced cyanophage fitness, increasing genome replication and progeny production during infection of *Synechococcus* WH8109. Furthermore, this thioredoxin was redox active and negatively impacted *Synechococcus* growth, indicating an underlying effect on host physiology. Deletion of *cve* showed that it increased cyanophage virulence, yet reduced progeny production. Despite this trade-off, it significantly enhanced overall cyanophage fitness, unveiling a host-type-specific virulence enhancer, that likely improves infection success and contributes to cyanobacterial mortality in the oceans. Thus, we named this gene *cve* for cyanophage virulence enhancer. Our findings suggest that virus-encoded thioredoxins are likely to provide an evolutionary advantage to a wide range of viruses and that at least some small, non-core viral dark matter genes are likely to be functionally important for phage infection and fitness, impacting microbial populations and organic matter recycling.

## Introduction

Cellular-like genes are prevalent in viruses (1–7) and are often acquired from their hosts over the course of evolution. These genes, known as auxiliary metabolic genes (AMGs), are common in viruses from diverse environments (3, 4, 8) and can reshape host metabolism to facilitate virus progeny production (5, 9–11). AMGs are particularly well-known in viruses that infect marine cyanobacteria (cyanophages) and include genes involved in photosynthesis (1, 12–14), carbon metabolism (15), nutrient utilization (16) and nucleotide metabolism (17).

Thioredoxins are ubiquitous redox enzymes found in cellular organisms from all domains of life (18, 19) with homologs in many virus genomes (9, 17, 20–26). In cells they are known to have multiple functions including regulating the activity of proteins based on the redox state of the cell, reduction of oxidized proteins as part of the cellular oxidative stress response, and the direct scavenging of reactive oxygen species (18, 19). In photosynthetic organisms such as cyanobacteria, thioredoxin is essential for growth (27), taking electrons generated during photosynthesis and passing them to transcription factors (28, 29) and metabolic enzymes (30), thus regulating their activity. The T7 bacteriophage uses the host’s thioredoxin to enhance T7 DNA polymerase activity in a redox independent manner (31–34). However, the function and evolutionary benefits of phage-encoded thioredoxins have not been investigated.

The genomes of phages from diverse environments contain a large number of small genes with no recognizable functional domains (35–39). These can account for 40-90% of viral open reading frames in environmental DNA and are often referred to as “viral dark matter”. In phage isolates the vast majority of these genes are non-core genes present in only a small subset of phages from a particular lineage (17, 23, 40–43). Their small size, lack of known domains and sporadic distribution have raised questions as to whether such phage genes have a functional role during infection and provide an evolutionary advantage to the viruses that carry them (44–46, 38).

Marine picocyanobacteria of the genera *Synechococcus* and *Prochlorococcus* are globally abundant phytoplankton (47, 48), estimated to be responsible for approximately 25% of ocean primary production (49). Cyanobacteria coexist in the oceans with a variety of dsDNA tailed cyanophages belonging to the class *Caudoviricetes* (50, 13, 51). The most abundant cyanophage groups are the lytic T4-like myoviruses (*Kyanoviridae*) and T7-like podoviruses (*Autographivirales*) (9, 52–56), although other cyanophages, including siphoviruses, TIM5-like myoviruses and others, are also known (57–61). T4-like and the T7-like cyanophages can infect large numbers of cyanobacteria (56, 62) and can therefore have a considerable impact on host populations(56). Cyanophage-induced lysis and intact infected cyanobacteria release dissolved organic matter which stimulates the growth of other microorganisms, contributing to the recycling of organic matter in the ocean (the viral shunt) (10, 63–65).

In this study, we explore the distribution of phage thioredoxin genes (*trx*) and a small gene of unknown function, *cve,* in phage isolates from various environments and assess their benefit to a T7-like cyanophage, Syn5, during infection of marine *Synechococus* WH8109. We found that thioredoxins are widespread in diverse phages infecting both ecologically and clinically relevant hosts whereas *cve* is restricted to marine cyanophages. While both genes increased the fitness of the Syn5 cyanophage, each gene contributed differently with the thioredoxin gene increasing phage genome replication and progeny production whereas the *cve* gene increased cyanophage virulence. Our findings demonstrate that phage-encoded thioredoxin and small genes of unknown function can have significant impacts on phage infection and fitness.

## Materials and methods

### Cyanobacterial growth and cyanophage propagation

*Synechococcus* sp. strain WH8109 (*Synechococcus* WH8109 from herein) was grown in artificial seawater medium (ASW) (66). Cultures were grown at a light intensity of 20-25 µmol photons m^-2^ sec^-1^ under a 14/10 day/night cycle and at a temperature of 21°C. Growth was inferred from daily measurements of chlorophyll *a* fluorescence (excitation at 440±20 nm, emission 680±20 nm) using a Synergy Mx microplate reader (BioTek, USA). *Synechococcus* WH8109 colonies and lawns were grown using the pour plate method with ASW medium supplemented with +SO_3_^2-^ and 0.28% low melting point agarose (67, 68) (Invitrogen, USA). To achieve high plating efficiencies, a ‘helper’ bacterium, *Alteromonas sp.* strain EZ55, was added whenever plating colonies to reduce oxidative stress (69).

Syn5 lysates were prepared by infecting exponentially growing *Synechococcus* WH8109 at a low MOI. After 24 hours the resulting lysate was filtered over a 0.22µm Millex GV syringe filter to remove cellular debris and stored at 4°C in glass tubes. The plaque assay was used to quantify the number of infective phages (plaque forming units - PFU). Filtered lysates were serially diluted, mixed with *Synechococcus* WH8109 cultures, pour-plated and grown to produce plaques as described above but without the addition of the ‘helper’ bacterium.

### Phylogenetics

Gene trees were built for thioredoxin using the NGPhylogeny.fr web server (70). Protein sequences were identified by sequence homology using PSI-BLAST and retrieved from NCBI. Thioredoxin homologs retained for our phylogenetic analyses all contained the canonical active site CGPC or CAPC (71), except for two T7-like cyanophages that had CQPC or CKPC.

Protein sequences were aligned using the MAFFT program and curated using TrimAI with a gap threshold of 0.3. Maximum likelihood and neighbor-joining analyses were performed using the PhyML+SMS and FastME programs and their topologies were compared using TreeViewer (72) to ensure overall similarity. To obtain confidence estimates for the inferred tree topology, 500 bootstrap resamplings were performed.

### Structural predictions and alignment

Protein structure models were predicted using AlphaFold3 (73). Predicted template modeling (pTM) score and interface predicted template modeling (ipTM) were provided by AlphaFold3 and used to assess confidence in the overall structure of complexes and relative positions of the subunits within (74). Structures were visualized using the ChimeraX software (75). Structural alignment was performed using the matchmaker function with default parameters.

### Vector construction and bacterial conjugation

Vectors for expression of a codon optimized Syn5 *trx* gene were ligated into the pBAD plasmid (Thermo Fisher Scientific) using the NcoI and PmeI restriction sites and electroporated into an *E. coli* strain lacking the endogenous thioredoxin gene (76) (Table S1). Transformed cells were selected for using 50 µg/mL ampicillin. For phage gene inactivation and expression of Syn5 *trx* in *Synechococcus* WH8109 inserts were generated using PCR overlap extension (77) and ligated into the pDS-proCAT (78) vector (Table S1) at BamHI and BsiWI restriction sites. Conjugation was performed following Brahamsha (79) with modifications as per Shitrit et al. (78) and with the *E. coli* S17.1 conjugative donor strain (80). Selection was done using ASW medium containing 1 µg/mL chloramphenicol.

### Thioredoxin activity assays

We used the thioredoxin fluorometric activity assay kit (Cayman Chemical) to measure the reduction of eosin-labeled insulin by thioredoxin in a cell lysate over time. Thioredoxin expression strains of *E. coli* were produced as described above. Protein expression was induced using 0.1% (v/v) L-arabinose (Sigma-Aldrich) and grown with vigorous shaking overnight at 37°C. See supplementary methods for more details.

### Phage gene inactivation

Phage gene inactivation mutants were generated using the REEP method described by Shitrit et al. (78). Briefly, a culture carrying a recombination template vector for gene inactivation was infected with the wild-type phage. A double homologous crossover between the phage genome and the inactivation template produces a deletion of the gene. The mutant phage was then isolated by enrichment and PCR screening and double plaque-purified. Full genome sequencing was performed to ensure an identical genetic background to that of the wild-type phage.

### Phage infection properties

Phage growth curves were used to assess phage infection properties. Cultures were infected at a multiplicity of infection (MOI) of 1. Samples were collected periodically, diluted 10-fold and filtered over a 0.22µm Millex GV syringe filter to remove cells and cellular debris and to collect extracellular phages in the filtrate. The number of infective phages was then quantified using the plaque assay.

Average burst sizes per cell and virulence were determined using population level assays (81). Cultures infected at an MOI of 1 were diluted 10^5^-fold after 60 minutes to prevent additional infections and were quantified using the infectious centers assay (82) which measures the difference in infectious units between unfiltered samples (infected cells and free phages) and 0.22µm filtered samples (free phages only). The number of infective phages produced was quantified 24 hours after infection. Burst size was calculated as the number of free infective phages produced per infected cell. Virulence was determined from the ratio of infectious centers to total number of living cells determined by plating colonies of a non-infected control.

### Competition assay

A competition assay was used to assess the contribution of *trx* and *cve* to competitive phage fitness (83). Equal concentrations of two phage strains were used to infect the same culture (MOI=10^-4^) over a 24 h period allowing for multiple infection cycles. Relative abundance of each strain was determined before and after competition by plating plaques and screening each plaque by PCR to determine its genotype (n=96). PCR was carried out using Taq PCR MasterMix (Tiangen) with 0.4 µM primers (Table S1).

### Phage DNA replication

Intracellular phage DNA replication was assessed by qPCR. Infection was performed at a cell concentration of ∼7.5*10^7^ cells/mL and an MOI of ∼3. Samples were subjected to quantitative DNA extraction (84, 85). Briefly, cells were collected on 25 mm, 0.2 μm polycarbonate filters (General Electric) by filtration, washed 3 times with ASW medium and once with preservation solution (10 mM Tris⋅HCl [pH 8], 100 mM EDTA, 0.5 M NaCl). Cells were removed from filters by immersion in 10 mM Tris⋅HCl (pH 8) and agitation in a bead-beater (Mini-BeadBeater, Biospec) for 2 minute (3450 revolutions/min) without beads. DNA was extracted by heat lysis at 95°C for 15 min. Phage DNA was quantified by real-time qPCR using primers for the DNA polymerase gene of Syn5 (Table S1). Reactions were carried out in a LightCycler 480 Real-Time PCR System (Roche) with a cycling program as previously described (85) with 500 nM desalted primers.

### Inducible expression system in *Synechococcus* WH8109

A theophylline-dependent riboswitch was used to test the effect of expression of different versions of the Syn5 thioredoxin on *Synechococcus* WH8109 growth. We used 0.1 mM theophylline to induce translation and monitored culture growth over 10 days. We compared induced to non-induced cultures to determine the effect of thioredoxin expression on bacterial growth and used a culture with an empty vector to control for the potential effect of theophylline. Five biological replicates were performed for each expression experiment. See Supplementary information for the adaptation of this method for marine *Synechococcus* WH8109 and for the determination of the appropriate conditions to use for induction of gene expression.

### Statistical analysis

All statistical analyses were performed using MathWorks Inc. MATLAB software, R2024a. For time-series data, we used repeated measures ANOVA, testing the assumption of sphericity (equality of variance of differences) using Mauchly’s test and applied the Greenhouse-Geisser correction to calculate the p-value if it was violated. For pairwise comparison of means, either the Student’s *t*-test (parametric) or Mann–Whitney *U* test (nonparametric) was used depending on the data distribution. Normality was tested using the Shapiro –Wilk test package (86). Latent period length was determined using the ischange MATLAB function to detect the initial rise in the log-transformed number of infective viruses. Log transformation was performed to increase the accuracy by reducing the impact of sampling variability.

## Results and discussion

In this study we first assess the prevalence of a putative thioredoxin gene (*trx*) and a small gene of unknown function (*cve*) in phage isolates from a variety of environments. We then investigate the functional role and fitness benefit of each gene for the Syn5 T7-like cyanophage during infection of marine *Synechococcus* WH8109 (52). The *cve* gene is directly downstream of, and overlaps with, the *trx* gene. Both genes are located amidst the DNA replication gene cluster of the Syn5 genome (20). We hypothesized that these genes augment phage fitness through enhancing phage DNA replication, both due to their position in the phage genome and the known use of host thioredoxin by the T7 coliphage (31–33).

### The thioredoxin gene is widespread in phages while *cve* is restricted to cyanophages

An investigation of the prevalence of the thioredoxin gene revealed that it is widespread in phages. Phages from diverse environments carry the gene, including those isolated from marine, terrestrial soil, freshwater, animal and human gut environments (Fig. 1A, Table S2). These phages also infect diverse host types of both ecological and clinical relevance, such as strains of *Synechococcus*, *Bacillus*, *Streptomyces, Acinetobacter, Pseudomonas, Escherichia coli, Burkholderia,* and others (Table S2). In addition, phages carrying the thioredoxin gene belong to a variety of taxonomic groups within the *Caudoviricetes* class (Table S2). In cyanophages thioredoxin is found in diverse T4-like cyanophages (25% of the 36 T4-like cyanophage genomes selected to represent genomic diversity (23)) and in one of three clades of T7-like cyanophages (clade A).

**Figure 1:**
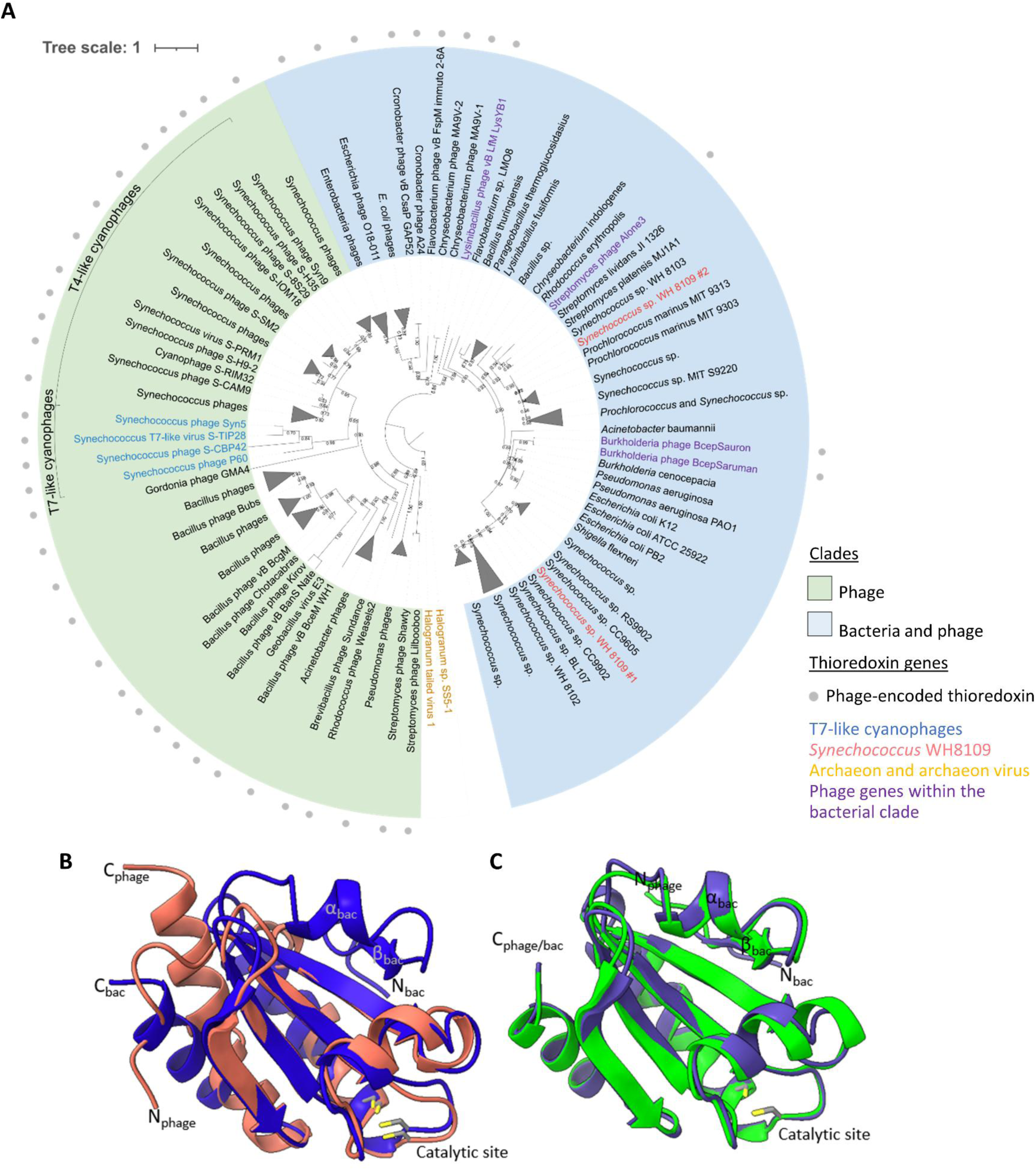
Gene tree and alignment of predicted structures of phage and bacterial thioredoxin. (**A**) Maximum-likelihood tree for thioredoxin, inferred from 100 amino acid positions. Bootstrap values equal to, or greater than, 50% are shown at branch nodes. The clade consisting of only phage thioredoxins is shaded in green, and the clade with both bacterial and phage thioredoxins is shaded in blue. Genes from phage isolates are indicated by a filled grey circle, T7-like cyanophages are shown in blue type, *Synechococcus* WH8109 genes in pink type, archaeon and archaeon virus in orange type and phage genes within the bacterial subclade are shown in purple type. Cyanophage clusters are indicated on the tree by brackets. (**B**) AlphaFold3 predicted structural alignment of *E. coli* K12 (blue) and Syn5 (red) thioredoxins, and (**C**) of *Burkholderia cenocepacia* (purple) and Burkholderia phage BcepSauron (green) thioredoxins. Carbon (grey) and sulfur (yellow) atoms of the cysteine residues of the catalytic sites are shown. Phage or bac indicate the location of phage or bacterial sections of the protein. N and C refer to the N or C-terminus of the proteins, α and β refer to α helixes and β strands.

Phylogenetic analysis shows that bacterial and phage thioredoxins cluster into two large clades. One of these clades consists exclusively of phage-encoded genes, including all those found in T4-like and T7-like cyanophages (Fig. 1A, green shaded). A second large clade contains all of the cellular copies as well as a distinct subclade that consists of phage thioredoxin genes (Fig. 1A, blue shaded). Assuming that thioredoxin is of cellular origin, these findings suggest that at least two independent transfer events gave rise to this distribution of thioredoxin in phages, followed by independent evolution and divergence from cellular homologs. Interestingly, there are four phage genes that cluster in close proximity to their respective hosts (Fig. 1A, purple typeface), suggesting independent and recent acquisitions from their bacterial hosts. Whether acquired recently or in the distant past, the widespread occurrence of thioredoxin genes in phages from different families and infecting a range of bacterial taxa from diverse environments suggests that thioredoxins confer an evolutionary advantage to diverse phages.

The predicted structure of phage thioredoxins is very similar to that of bacterial thioredoxins (Fig. 1B). Sequence, position and predicted conformation of the catalytic site (CXPC) is conserved among bacterial and phage thioredoxins (Fig. 1B, Figure S1). However, thioredoxin in the two phage clusters differ at the N-terminus, lacking approximately 20 amino acids found in cellular copies (Fig. 1B, Fig. S1), suggesting that the N-terminus was either lost by phages twice or gained by bacteria. In contrast, the phage thioredoxins that cluster with their hosts still contain the bacterial-like thioredoxin N-terminus (Fig. 1C). The overall structural similarity yet differences at the N-terminus, raises the question as to whether phage thioredoxins function similarly or differently to bacterial thioredoxins.

Next, we assessed the distribution of the *cve* gene in phages. In contrast to thioredoxin, we found that this gene is restricted to cyanophages isolated from the marine environment. It is predominantly found in T4-like cyanophages (69.4% of 36 representative T4-like cyanophage genomes (23)), and is also present in some T7-like cyanophages, TIM5-like phages and siphoviruses that infect marine cyanobacteria (Table S3). Interestingly, we found a homolog of this gene in a putative complete PSS2-like cyanopsiphovirus prophage that we detected in the genome of *Synechococcus* sp. SYN20 (see Supplementary text). The restricted presence of *cve* to marine cyanophages, while occurring in different cyanophage families, suggests specific importance for infection of these cyanobacteria.

A small subset of cyanophages encode both the thioredoxin and *cve* genes (Table S3B, 17%). In two of the four T7-like cyanophages with both genes, including Syn5, the start codon of the *cve* gene overlaps with the stop codon of the thioredoxin gene (Table S3A). This is known to occur in viruses and often results in dependence of translation of the second protein on the translation of the first protein, known as translational coupling.

### Thioredoxin and Cve are translationally coupled and increase phage fitness

The prevalence of thioredoxin and *cve* in cyanophages raises the possibility that these genes contribute to phage fitness. To test this hypothesis, we used our recently developed genetic manipulation system, REEP (78), to delete each of these genes from the Syn5 phage. However, because of the overlap between the two genes, it was important to first assess whether they are translationally coupled, since this would cause the Cve protein to be missing in the *trx* deletion mutant (Δ*trx*). To test this, we measured peptide levels of the two proteins in mutant and wild-type phages (see Supplementary methods). In line with our expectation, neither thioredoxin nor Cve peptides were detected in the Δ*trx* mutant phage while both were present in the wild-type phage (Fig. S2A). Transcripts of the *cve* gene were at a similar level in both phages (Fig. S2B). These findings indicate that translation of Cve is coupled to that of thioredoxin. Translational coupling was restricted to these two proteins since similar levels were found of the protein coded directly downstream of *cve,* the DNA polymerase (Fig. S2).

Such translational coupling could be indicative of functional coupling and/or of genome streamlining (87). Since both genes are not consistently present in cyanophages and overlapping start and stop codons are found in only two phages, we consider genome streamlining to be the more likely explanation in this case. The consequence of such translational coupling for our study is that the Δ*trx* cyanophage mutant is a functional double mutant lacking both the thioredoxin and the Cve proteins. We will thus refer to this mutant as Δ*trx*/Cve^S^, with the “s” referring to silencing.

We next turned to assessing the relevance of these two genes for cyanophage fitness, both together and individually. For this, direct competition experiments were carried out in the same flask over multiple infection cycles. Direct competition between the *Δtrx*/Cve^S^ mutant and the wild-type phage revealed that the wild-type outcompeted the mutant, producing ∼6-fold more phages than the mutant (Fig. 2A, p=0.00012, n=4). This indicates that together these genes contribute significantly to competitive phage fitness.

**Figure 2:**
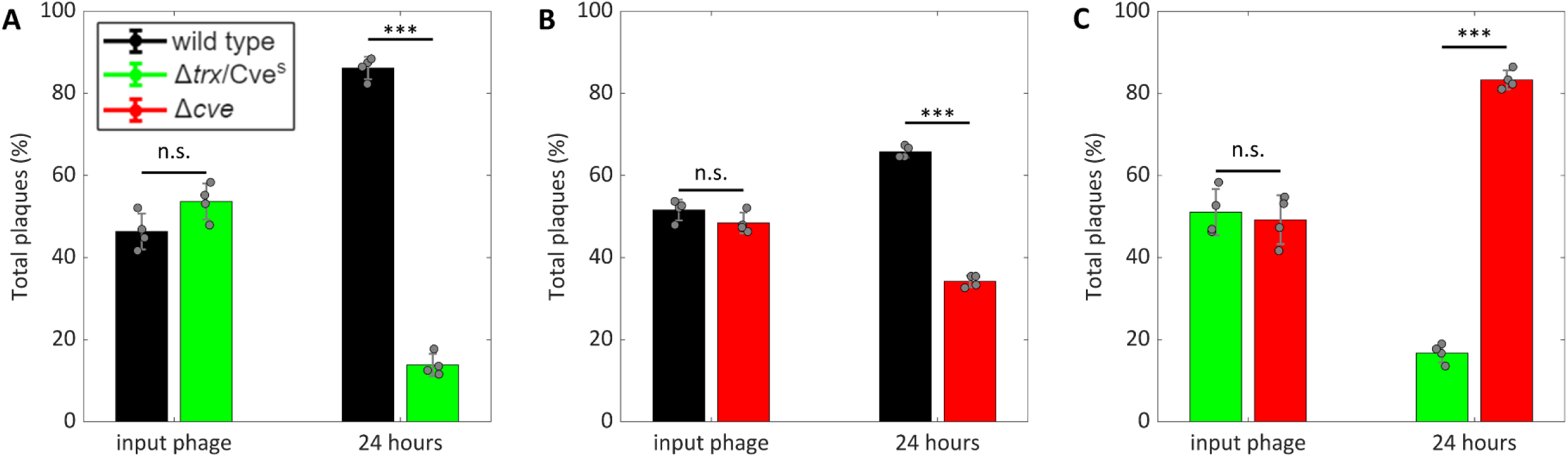
Direct competition between wild type and mutant Syn5 strains. A comparison of the relative abundance of the Syn5 wild type and the *Δtrx*/Cve^S^ mutant (**A**), the wild type and the Δ*cve* mutant (**B**) and the Δ*cve* and *Δtrx*/Cve^S^ mutants (**C**) infecting the same host cultures before (input phage) and after (24 hours) competition. The wild type is shown in black, the *Δtrx*/Cve^S^ mutant in green and the Δ*cve* mutant in red. Averages and standard deviations of 4 biological replicates. P-values of paired, two tailed student t-test are shown. *** - p-value<0.001. n.s. - not significant.

Next, we assessed the contribution of each gene individually. Competition between the *Δcve* mutant and the wild-type phage revealed that ∼2-fold more of the wild-type phage was produced than the *Δcve* mutant (Fig. 2B, p=0.00021, n=4), indicating that Cve contributes to competitive phage fitness independently of thioredoxin. We then compared the *Δtrx*/Cve^S^ and *Δcve* mutants to assess the effect of thioredoxin alone on phage fitness. We found ∼5-fold more of the *Δcve* mutant than the *Δtrx*/Cve^S^ mutant (Fig. 2C, p=9.2*10^-5^, n=4). While *trx* and *cve* both have individual contributions to competitive phage fitness, the effect of *trx* was more substantial as seen from the greater fold difference between the *Δcve* and the *Δtrx*/Cve^S^ mutants than between the wild-type and the *Δcve* mutant (Fig. 2B,C).

### Thioredoxin and Cve differentially alter cyanophage infection properties

Phage fitness is impacted by different infection properties. These include the rate of adsorption, the length of time needed to produce new phage progeny (the latent period), the average number of infective progeny produced per cell (the burst-size) and the percentage of cells lysed during infection (the virulence) (88). We set out to elucidate the effects of *trx* and *cve* on these cyanophage infection properties using the same wild-type and mutant comparisons as described above.

To test the effect of the genes on the rate of adsorption and the length of the latent period we used phage growth curve experiments. We found that neither gene altered the rate of adsorption nor the length of the latent period, remaining at 1.5 h (Fig. 3A). These experiments also provide a measure of phage production over time which integrates across several infection properties within a single infection cycle (e.g. latent period, burst size and virulence). Phage production over time was lower in the *Δtrx*/Cve^S^ mutant compared to the wild-type phage (Fig. 3B, p=0.015, n=4) indicating that both genes together increased progeny production. In fact, each gene individually contributed to phage production over time with greater production in the wild-type relative to the *Δcve* mutant (Fig. 3C, p=6.6*10^-5^, n=4) and in the *Δcve* mutant relative to the *Δtrx*/Cve^S^ mutant (Fig. 3D, p=0.026, n=4).

**Figure 3:**
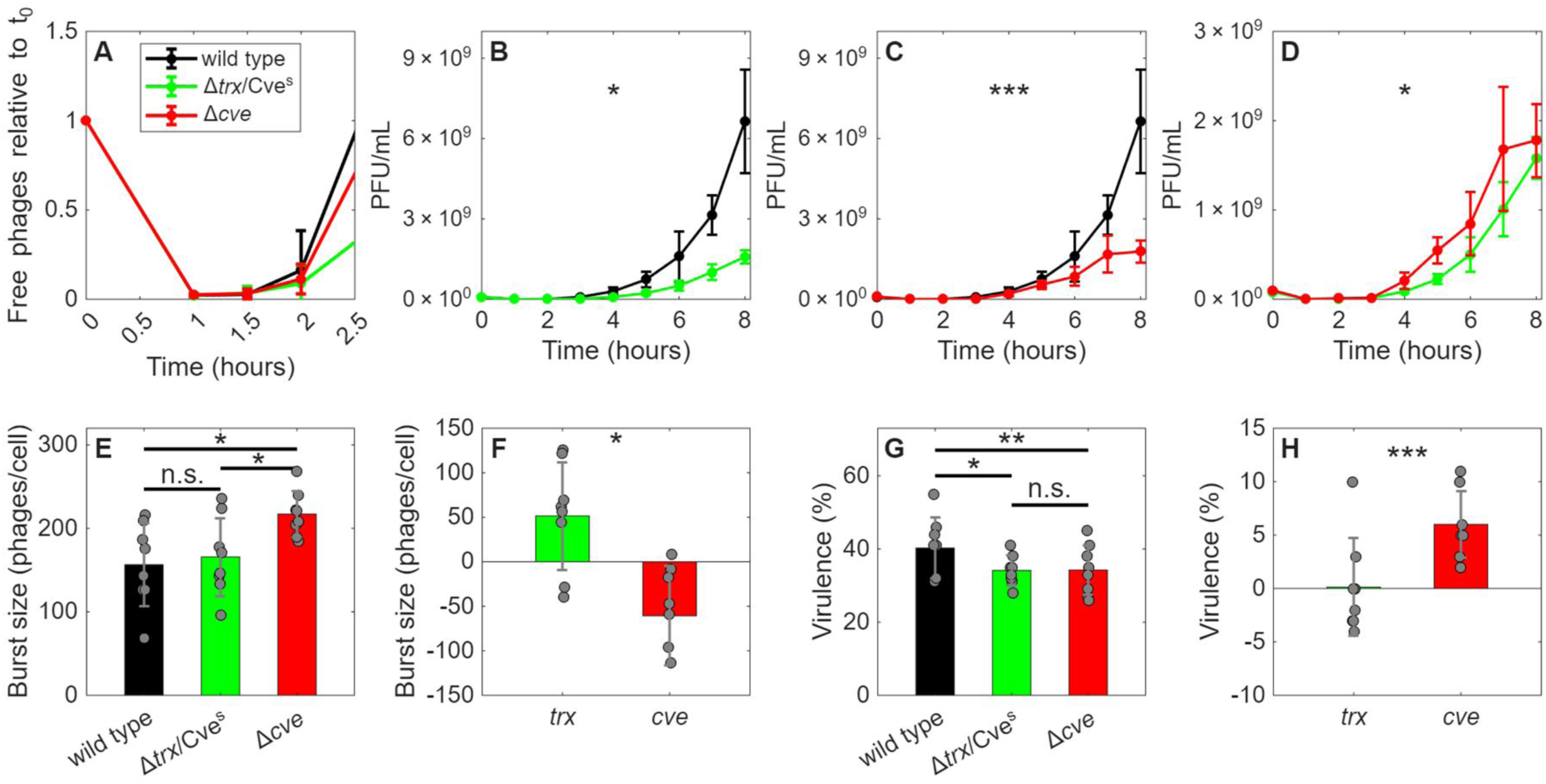
Effects of Syn5 *trx* and *cve* mutants on phage infection properties. Comparison of the latent period between the wild type and mutant phages (**A**). Comparison of phage progeny production over time between the Syn5 wild type and the *Δtrx*/Cve^S^ mutant (**B**), the wild type and the Δ*cve* mutant (**C**) and the Δ*cve* and the *Δtrx*/Cve^S^ mutants (**D**). Burst size (**E**), virulence (**G**) of the Syn5 wild type (black) and the *Δtrx*/Cve^S^ (green) and Δ*cve* (red) mutant strains and the contribution of *trx* and *cve* to burst size (**F**) and virulence (**H**). The contribution of *cve* to virulence or burst size was calculated as the differences between the wild type and Δ*cve* mutant. The distinct contribution of *trx* was calculated as the difference between the wild type and *Δtrx*/Cve^S^ after subtracting the contribution of *cve*. For A-D, average and standard deviation of 3 biological replicates for latent period comparison and 4 biological replicates for progeny production over time. P-values of significantly different numbers of plaque forming units (PFU) produced over time (repeated measures ANOVA), are shown. * - p-value<0.05, *** - p-value<0.001. For E-H, average and standard deviation of 8 biological replicates. P-values of significantly different virulence or burst size (Wilcoxon signed rank test or paired t-test) are represented by: * - p-value<0.05, ** - p-value<0.01 *** - p-value<0.001, n.s. - not significant.

Next, we tested the effect of *trx* and *cve* on the burst size. The burst size of the wild-type phage averaged 156.5±49.8 phages·cell^-1^ (Fig. 3E). Intriguingly, the burst size of the *Δcve* mutant was significantly higher (217.2±27.4 phages·cell^-1^) (Fig. 3E, p=0.016, n=8), indicating that *cve* reduces the phage burst size by 60.7±56 phages·cell^-1^ (Fig. 3F). This corresponds to a decline of approximately 28% compared to the wild-type phage. However, the burst size of *Δtrx*/Cve^S^ and the wild-type phage were similar (Fig. 3E, p=0.55, n=8) suggesting that the negative effect of *cve* is mitigated by a positive effect of the *trx* gene. Indeed, a direct comparison between the *Δtrx*/Cve^S^ and Δ*cve* mutants showed that the lack of the thioredoxin gene resulted in a lower burst size by 51.5±60.5 phages·cell^-1^ (Fig. 3F). Thus, the *trx* gene results in an increase in burst size of approximately 33% relative to the wild-type phage. These results indicate that *cve* decreases, while *trx* increases, the burst size by a comparable amount, each canceling the effect of the other gene.

We next assessed the virulence of our phages. Wild-type phage virulence was 40.3±8.4% (mean ± SD), indicating that 40% of the cells in the culture were lysed (Fig. 3G). In comparison, *Δcve* mutant virulence was 34.1±6.8% (Fig. 3G), lower than that of the wild-type by 6% (Fig. 3G, B, p=0.0078). When normalizing the number of lysed cells to the wild-type phage, this indicates that *cve* increased virulence by ∼15%. However, no difference in virulence was observed between the Δ*cve* and *Δtrx*/Cve^S^ strains (Fig. 3G, H, p=0.69), indicating that *trx* does not contribute to phage virulence.

Our results indicate that *trx* and *cve* each have distinct, independent contributions to phage fitness despite begin translationally coupled. The *trx* gene positively affects burst size and is likely to be the infection property responsible for the increased progeny production over time and enhanced cyanophage fitness (Fig. 2C, 3D-F). In comparison, *cve* enhances phage virulence at the expense of burst size (Fig. 3E-H), yet still increases progeny production over time and phage fitness (Fig. 2B, 3C). This indicates that virulence is particularly important for phage fitness, with the increase outweighing the negative effect on burst size. As such, we named the gene *cve* for cyanophage virulence enhancer.

The cyanophage-specific *cve* gene is likely to have a direct impact on ocean ecosystems. First, the higher virulence attributable to the gene, together with its widespread occurrence in marine cyanophages, could have a detrimental impact on cyanobacterial population sizes (88). Furthermore, the higher number of cells lysed by *cve*-carrying cyanophages would increase the amount of overall dissolved organic matter released from infected cyanobacteria after lysis that becomes available to other microorganisms (63, 65). As such, this gene is likely to influence the marine carbon cycle by contributing to the viral shunt.

### Cyanophage thioredoxin increases phage genome replication

To test our hypothesis that *trx* and/or the *cve* contribute to phage genome replication we measured intracellular phage DNA levels during infection. We found that phage DNA was lower during infection with the *Δtrx*/Cve^S^ mutant compared to infection with either the wild-type or the Δ*cve* mutant phages (Fig. 4A, B, p=0.013 and 0.01 respectively, n=4). However, there was no difference in genome replication between the wild-type and Δ*cve* mutant phages (Fig. 4C, p=0.27, n=4). These findings indicate that the *trx* gene, but not the *cve* gene, is responsible for increased phage genome replication.

**Figure 4:**
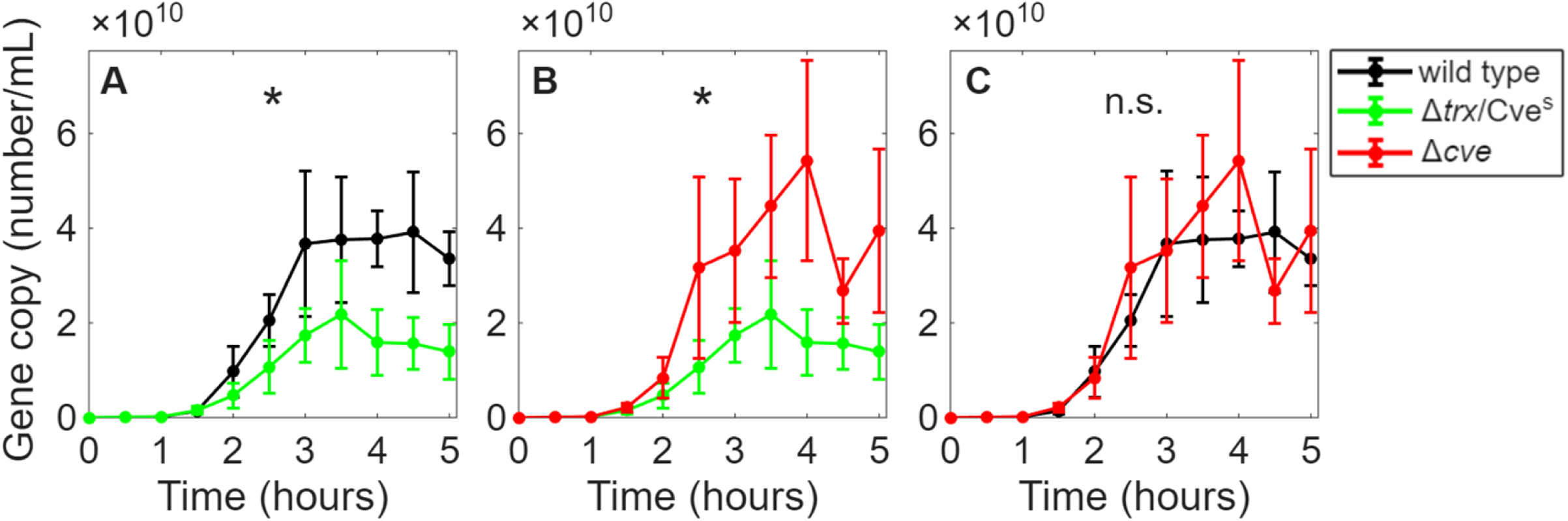
The effect of *trx* and *cve* on phage DNA replication. A comparison of intracellular DNA replication between the Syn5 wild type and the *Δtrx*/Cve^S^ mutant (**A**), the Δ*cve* and *Δtrx*/Cve^S^ mutants (**B**) and the wild type and the Δ*cve* mutant (**C**). The wild type is shown in black, the *Δtrx*/Cve^S^ mutant in green and the Δ*cve* mutant in red. Average and standard deviation of 4 biological replicates. P-values of significantly different intracellular phage DNA copy numbers over time (repeated measures ANOVA), are shown. * p-value<0.05, n.s. - not significant.

The phage-encoded thioredoxin could enhance phage DNA replication through two different mechanisms (20, 22). One possibility is through its redox activity, activating the phage’s ribonucleotide reductase for conversion of host ribonucleotides to deoxyribonucleotides (20, 22) and increasing the availability of nucleotide substrates for phage genome replication. Alternatively, the mode of function of the cyanophage thioredoxin may be independent of its redox activity through direct binding to its DNA polymerase, increasing processivity (20) and enabling more efficient phage-genome replication. This latter option is the mechanism in which the T7 bacteriophage uses the *E. coli* cellular thioredoxin (31–33). To begin exploring these possible mechanisms, we first determined whether the Syn5 thioredoxin has redox activity. For this we employed an in vitro redox activity assay and found that the Syn5 cyanophage carries an enzymatically functional thioredoxin (Fig. 5A, p=8.66*10^-286^, n=4) and that the catalytic site, which is identical to that of cellular thioredoxins, is essential for this activity (Fig. 5B, C, p=0.52 and 0.5, n=3, see supplementary text for more details).

**Figure 5:**
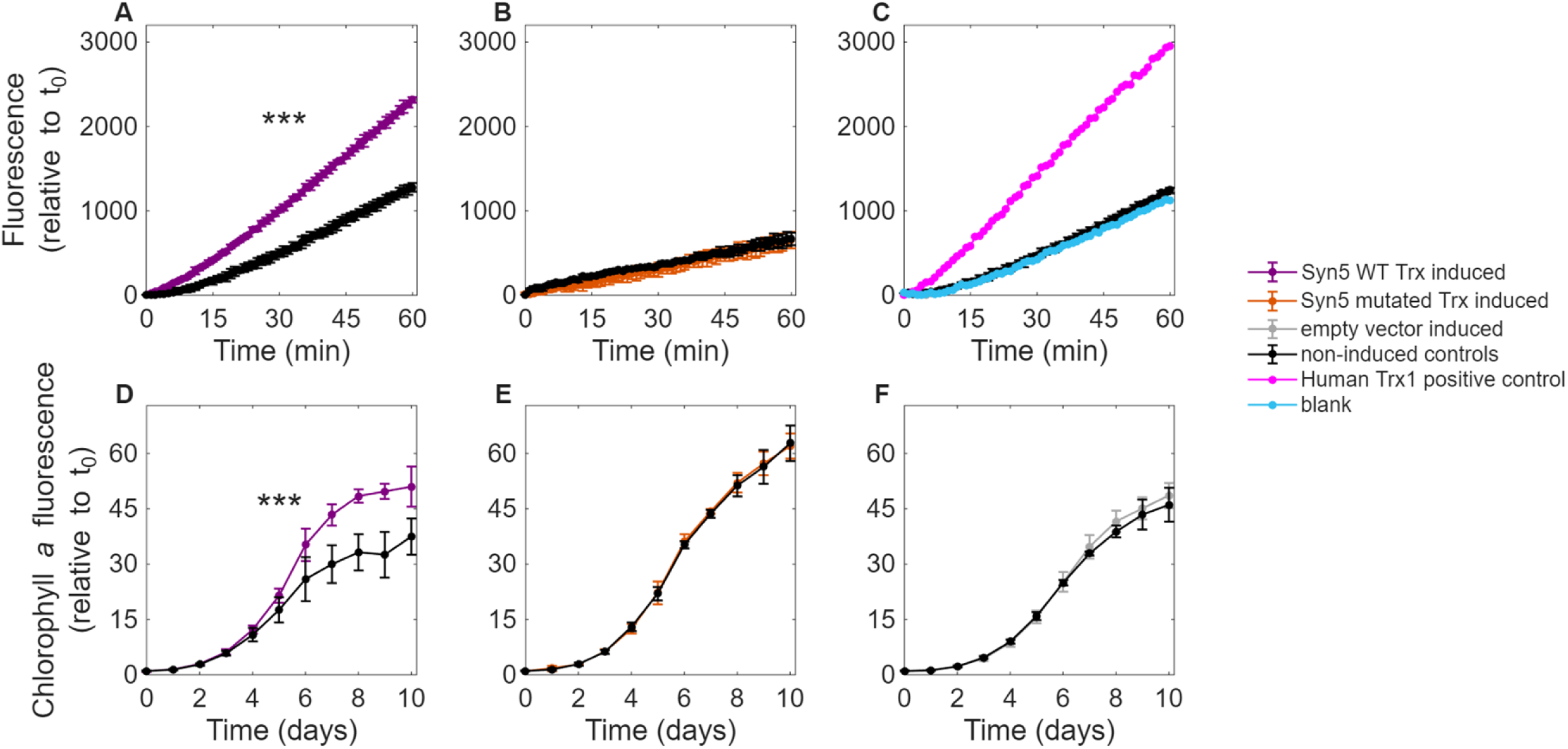
Comparison of the redox activity of the Syn5 phage wild type and mutated thioredoxin and their effect on *Synechococcus* growth. **(A-C)** Redox activity of the Syn5 wild type thioredoxin **(A)** and the Syn5 thioredoxin with a mutated catalytic site **(B)** when expressed from inducible vectors in an *E. coli* strain lacking its own thioredoxin gene. Positive control (human thioredoxin protein in pink) and negative control lacking the gene (empty vector) **(C)**. Expression of the gene was induced by adding arabinose (colored lines) and compared to non-induced controls (black lines). Note that in the empty vector control the induced (grey) and non-induced (black) lines fall on top of each other. Cell lysates were assayed for their ability to reduce eosin-labeled insulin. Average and standard deviation of 3 biological replicates for mutated thioredoxin and 4 biological replicates for WT thioredoxin and the empty vector. **(D-F)** Growth curves of *Synechococcus* WH8109 expressing the Syn5 wild type thioredoxin **(D)**, the Syn5 thioredoxin with a mutated catalytic site **(E)** and the empty vector **(F)**. Expression was induced by adding theophylline (colored lines) and compared to non-induced controls (black lines). Average and standard deviation of 5 biological replicates. The effect of the inducer was tested using the repeated measures ANOVA. *** p-value<0.001.

To assess the possibility of redox-independent use of cyanophage thioredoxin through binding to its DNA polymerase we predicted their structural interaction using AlphaFold3 (73). As a control, we compared the modeled complex with the experimentally determined complex of the T7 DNA polymerase and the *E. coli* thioredoxin and found that they are nearly identical (Fig. 6A, S3, supplementary text) providing confidence in this comparative approach (89). In comparison, no confident structural interaction was predicted for the Syn5 cyanophage thioredoxin and its DNA polymerase. Rather, only a low confidence prediction of binding between the two proteins was generated and this was to a different site on the cyanophage DNA polymerase (Fig. 6B). These findings suggest that it is highly improbable that the cyanophage thioredoxin functions as a processivity factor for the cyanophage DNA polymerase. While we lack direct evidence at this stage, these structural predictions together with thioredoxin’s redox activity suggest that the more likely mode of action of the Syn5 cyanophage thioredoxin in enhancing phage genome replication is through activation of phage ribonucleotide reductase.

**Figure 6:**
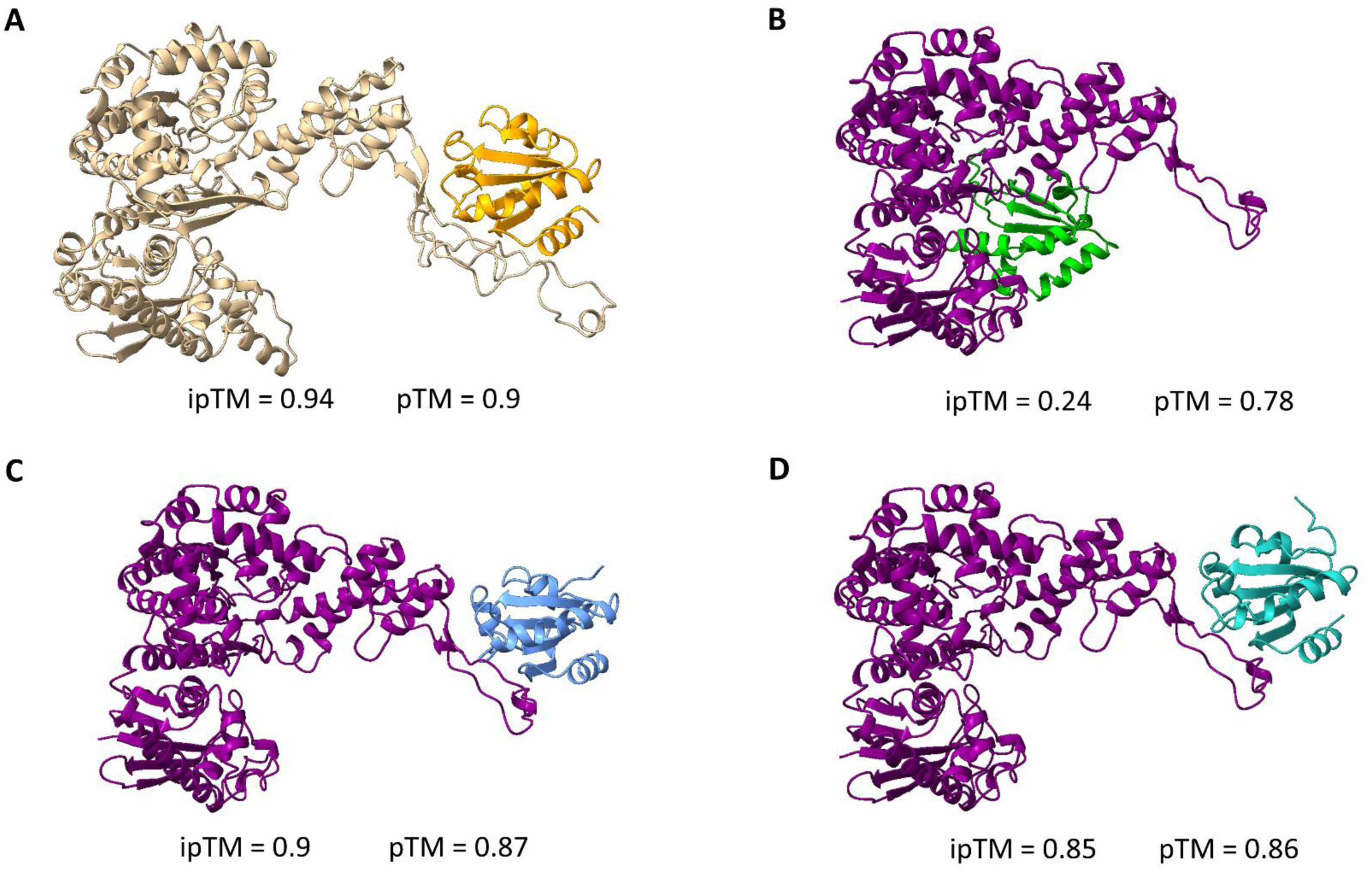
AlphaFold3 predicted structures of phage DNA polymerase interacting with different thioredoxins. (**A**) Predicted fold of T7 DNA polymerase (beige) with *E. coli* K12 thioredoxin (orange), (**B**) Syn5 DNA polymerase (purple) with Syn5 thioredoxin (green), (**C**) Syn5 DNA polymerase (purple) with *Synechococcus* WH8109 thioredoxin 1 (blue) and (**D**) Syn5 DNA polymerase (purple) with *Synechococcus* WH8109 thioredoxin 2 (teal). pTM - predicted template modelling scores of the proteins in the complex, ipTM - interface predicted template modelling scores of complex formation and the relative positions of the proteins. Higher scores indicate higher confidence prediction. ipTM values above 0.8 represent confident high-quality predictions whereas values below 0.6 are considered unreliable (74). GenBank accession numbers are AHF64381.1 and AHF63033.1 for *Synechococcus* WH8109 thioredoxin 1 and 2 respectively.

Interestingly, we found that the two thioredoxins coded by *Synechococcus* WH8109 are predicted to bind the Syn5 DNA polymerase with high confidence (Fig. 6C-D) in the same manner and position as the *E. coli* thioredoxin and T7 DNA polymerase (Fig. 6). This raises the possibility that Syn5 uses both its own thioredoxin and that of its host to enhance DNA replication. Therefore, we speculate that Syn5 and other diverse phages have two different complementary mechanisms to enhance phage genome replication; through the carriage and redox activity of their own thioredoxin activating ribonucleotide reductase and the exploitation of their host’s thioredoxin functioning as a DNA polymerase processivity factor.

### Cyanophage thioredoxin is toxic to the host

Cellular thioredoxins mediate many important redox-related activities and are essential in cyanobacteria (27–30). We tested for potential effects of the Syn5 cyanophage thioredoxin on the host using a theophylline-dependent riboswitch inducible translation system (90, 91) adapted for use in marine *Synechococcus* WH8109 (Supplementary information). We found that expression of the catalytically active thioredoxin negatively affected host growth (Fig. 5D) but that growth was not affected by a catalytically inactive thioredoxin (Fig. 5E). Thus, the Syn5 thioredoxin is detrimental to the *Synechococcus* host and this effect is mediated through its catalytic redox activity.

Since thioredoxins play many roles in photosynthetic organisms, the cyanophage thioredoxin could affect host growth through one or more multiple pathways. These include the combined activity of the phage thioredoxin and ribonucleotide reductase, resulting in the depletion of ribonucleotides needed for host RNA synthesis and thus growth. Alternatively, the effect could be due to reduction of host proteins that interferes with normal cellular redox functions (28, 29), or the redirection of redox power away from host processes such as carbon fixation (10, 15). Irrespective of the mode of impact of the phage thioredoxin on host physiology, our findings indicate that the cyanophage thioredoxin is both advantageous for phage fitness and detrimental to the host.

The phage thioredoxin gene is likely to have important downstream ecological consequences. Since thioredoxin impacts host physiology it is expected to have substantial metabolic effects on the infected cell. Furthermore, enhancement of phage genome replication is likely driven through changes in the quantity and composition of the nucleotide pool in the cell, regardless of the mode of action of the phage thioredoxin. Such changes would affect the composition and flux of the metabolites secreted from the cell during infection and after cell lysis, altering the dissolved organic matter available to heterotrophic bacteria (10, 65, 92) which would influence microbial community composition (64, 93, 94).

## Conclusions

Here we investigated two accessory cyanophage genes. Even though neither are core nor essential, both *cve* and *trx* enhance phage fitness, doing so via distinct mechanisms. *cve* increases phage virulence resulting in the successful infection and mortality of more host cells, and *trx* boosts phage progeny production presumably through the increase in genome replication. As such, in nature *cve* is more likely to influence host population sizes while *trx* would have a greater impact on phage population sizes. These two genes are also likely to affect the recycling of organic matter fixed by marine cyanobacteria in distinct ways, *cve* by increasing the amount of organic matter released through greater cell mortality, and *trx* by altering its composition. These resulting changes in the quantity and composition of released organic matter are thus likely to influence virus-induced carbon cycling in the oceans.

The distinct distribution patterns of the *trx* and *cve* genes are intriguing. The widespread distribution of *trx*, together with its clear fitness benefit (Fig. 2), suggests that this gene provides an evolutionary advantage to many dsDNA viruses infecting a wide range of distinct host types from a variety of environments. In contrast, the restricted distribution of the *cve* gene to phages infecting marine cyanobacteria, while from different cyanophage families, suggests a unique evolutionary advantage when infecting these particular hosts. This also raises the possibility that factors modulating virulence require tailoring for specific host types, and unveils a host-type-specific virulence enhancer in viruses.

Genomics has revolutionized viral ecology, unveiling the presence of auxiliary metabolic genes (1, 4, 7, 17) and the immense diversity of viral dark matter present in viral genomes (35–39). Yet, at best, we can propose potential functions when an identifiable domain is present, whereas in most cases insufficient information is available to even formulate an initial hypothesis. Thus, one of the greatest challenges facing the field is unravelling the functional role and ecological relevance of such genes. Our findings, while only a drop in the bucket, show that such auxiliary metabolic genes and non-core, non-essential genes can have important functions for the viruses that carry them, impacting their fitness, shaping their evolutionary trajectory and likely influencing ecological interactions.

## Supporting information

Table S2 - phage host pair and taxonomy

Table S3 - extensive and representative thioredoxin

## Acknowledgments

We thank Idan Yelin and Shirly Larom for their suggestions and discussion on the thioredoxin activity assay; David Keho for his suggestions and guidance with the GusA assay; Udi Qimron for the idea to try and complement T7 with Syn5 and host thioredoxin; Guy Horev for discussions on statistics; Oded Béjà and Lindell lab members for their support and discussions. Protein analyses were carried out at the Technion Smoler Proteomics Center. This research was funded by the European Research Council Consolidator Grant 646868, the Simons Foundation Life Sciences Grants 735081 and SFI-LS-PROJECT-12024 to DL.

