## Supplementary information for "Cyanophage thioredoxin and virulence enhancer genes differentially contribute to phage fitness and infection of marine *Synechococcus*"

### Supplementary results

#### Optimization of inducible expression system for *Synechococcus* WH8109

We adapted an inducible expression system at the stage of translation for marine *Synechococcus* following that used previously for diverse freshwater cyanobacteria (1, 2). This system utilizes a theophylline-dependent riboswitch of type E* (also known as F) that was engineered to consist of the *Prochlorococcus* MED4 *rnpB* promoter constitutively transcribing the riboswitch and the downstream gene of interest in the pDS-proCAT self-replicating plasmid (3).

To evaluate timing and changes in the level of gene expression from the riboswitch system, we employed the β-glucuronidase (GusA) reporter gene (4) in an assay previously described for *Fremyella diplosiphon* UTEX 481 (5, 6) (see Supplementary methods). We tested GusA activity between 0.5 to 72.5 hours after addition of theophylline at concentrations ranging from 0 to 0.3 mM (Fig. S4). GusA expression reached a plateau at 48.5 hours after theophylline addition at all concentration (Fig. S4). GusA expression levels increased with higher theophylline concentration up to 0.1 mM and were similarly high for theophylline concentrations ranging from 0.1 and 0.3 mM (Fig. S4). These results indicate that for marine *Synechococcus* WH8109 this expression system is best used at a minimal concentration of 0.1 mM theophylline and that induction be allowed to continue for a minimum of 48 h to achieve maximal expression levels.

#### Putative prophage in the genome of *Synechococcus* sp. SYN20

During our investigation into the prevalence of *cve* in the non-redundant database we found this gene to be limited to cyanophages except for *Synechoccucus* sp. SYN20. We hypothesized that this *Synechococcus* strain may have a prophage carrying the gene. To investigate this, we used PHASTEST (PHAge Search Tool with Enhanced Sequence Translation) (7) to look for prophages in the genome of this *Synechoccucus* strain. We found a putative PSS2-like cyanosiphoviruses prophage region within the genome of *Synechoccucus* sp. SYN20. The *cve* homolog was found within this region of the genome. Therefore, this result indicates that *cve* homologs are indeed only found in cyanophages.

**Measuring the catalytic activity of cyanophage thioredoxin in-vitro**

To begin addressing whether the phage thioredoxin is likely to function through its redox activity or independently of it, we first determined whether the Syn5 thioredoxin has redox activity. For this we expressed the Syn5 thioredoxin gene under an inducible promoter in an *E. coli* strain lacking its own thioredoxin gene and employed an in vitro redox activity assay. Induction of expression of the phage gene resulted in the reduction of the assay substrate, as seen from the increase in fluorescence over time relative to the non-induced control (Fig. 5A, $p=8.66*10^{-286}$, n=4). Modification of the two cysteines in the putative catalytic site (CGPC) to two serines (following Huber et al. (8)) abolished the redox activity of the phage thioredoxin, with no differences in fluorescence levels over time relative to the non-induced control as observed for the empty vector (Fig. 5B, C, p=0.52 and 0.5, n=3). These results indicate that the Syn5 cyanophage carries an enzymatically functional thioredoxin capable of substrate reduction and that the catalytic site, known for cellular thioredoxins, is essential for this activity.

### Supplementary methods

**Thioredoxin activity assay**

We used the thioredoxin fluorometric activity assay kit (Cayman Chemical) to measure the reduction of eosin-labeled insulin by thioredoxin in a cell lysate over time. Thioredoxin expression strains of *E. coli* expressing the Syn5 phage thioredoxin with wild type or mutated catalytic site (mutated, CGPC to SGPS), were produced as described in the main text. Protein expression was induced using 0.1% (v/v) L-arabinose (Sigma-Aldrich) and grown with vigorous shaking overnight at 37°C.

*E. coli* cells expressing Syn5 thioredoxin were harvested by centrifugation for 5 min at 5467g and 4°C, washed twice with ice-cold 100 mM Tris-HCl, 1 mM EDTA pH7.5 supplemented with protease inhibitor (Cayman Chemical) and resuspended in a tube containing ice-cold, acid-washed glass beads (Sigma-Aldrich). The culture was lysed using a bead-beater (Mini-BeadBeater, Biospec) for 1 min (3450 revolutions/min) followed by cooling on ice for 1 min. Cellular debris were precipitated by centrifugation for 3 min at 20817g and 4°C and the supernatant was used immediately to measure thioredoxin activity.

Thioredoxin activity in the cell lysate was tested by measuring the reduction of eosin-labeled insulin. The reduction of disulfide bonds in the labeled insulin by thioredoxin (9) releases eosin and results in an increase in eosin fluorescence (excitation at 520 nm and emission at 560 nm). This kit includes thioredoxin reductase and NADPH to recycle thioredoxin which is oxidized in the process of this reaction. Equal amounts of protein were used in the activity assay with protein concentrations being measured using the Pierce™ BCA Protein Assay Kit (Thermo Scientific).

**GusA quantitative activity assay**

We induced GusA expression in *Synechococcus* WH8109 using different concentrations of theophylline and measured GusA catalytic activity at several time points up to 72.5 h after induction (Fig. S4). Translation was induced by adding theophylline dissolved in ASW medium to an exponentially growing culture containing the expression plasmid. Cells were harvested by centrifugation at room temperature at 5467g for 5 min. Cell pellets were resuspended in ice-cold GUS assay buffer (1 mM EDTA, 50 mM $NaPO_4$ pH7) containing 0.001% SDS and 6.25 µg/mL chloramphenicol, then lysed by bead-beating and precipitated by centrifugation as described for the thioredoxin activity assay. Cell lysates were mixed with the GUS assay buffer containing 1.125 mM 4-Nirophenyl β-D-glucuronide (PNPG, Sigma-Aldrich). The degradation of PNPG by GusA, produces a yellow pigment which was measured by absorbance at 410 nm every 2 min for 30 min using a Synergy Mx microplate reader (BioTek, Winooski, CA, USA). The linear part of the slope of the resulting curve of absorbance over time was used to calculate GusA activity. Protein concentration in the cell lysates was measured using the BCA protein determination kit. GusA activity was quantified as nanomole of PNG produced per mg of protein per minute. Four biological replicates were performed for each theophylline concentration.

**Phage gene and protein levels**

Transcription and translation of specific genes were determined for the wild-type and mutant Syn5 phage strains. Infection was performed at a cell concentration of ~$7.5*10^7$ cells/mL and an

MOI of ~3. Cells were collected by centrifugation 60 min after infection at 15,000g for 2 min at 4°C for transcription and 5407g for 5 minutes at 21°C for translation. Cell pellets were flash frozen in liquid nitrogen and stored at -80°C prior to extraction and analysis. RNA and proteins were extracted as previously described by Zborowsky & Lindell (10) .

For the extraction of RNA, cell pellets were thawed, resuspended in 10 mM Tris·HCl (pH 8) with 100 units of RNase inhibitor (Applied Biosystems) and treated with 15000 units of lysozyme (Sigma-Aldrich) for 30 min at 37°C to lyse the cells. RNA was isolated using the Quick RNA Mini Prep Kit (Zymo) and residual DNA was removed using 2 units of TURBO DNase™ (Invitrogen). Reverse transcription (RT) was conducted using LunaScript™ RT SuperMix Kit (New England Biolabs). Random hexamer primers (6 µM) were annealed to RNA at 25°C for 2 min, followed by cDNA synthesis at 55°C for 10 min and heat inactivation at 95°C for 1 min. The cyanobacterial *rnpB* gene was used as a positive control for RT for all samples. No-RT controls were performed on all samples to ensure that reported levels were not from residual phage DNA. Transcript abundance was quantified using real-time qPCR carried out in a LightCycler 480 Real-Time PCR System (Roche) with a cycling program as previously described (10) with 500 nM desalted primers (Table S1). RT samples were stored at −20°C prior to qPCR.

For phage protein levels, proteins were trypsin digested in 1% SDC and 50 mM ammonium bicarbonate and peptides were resolved by reverse-phase HPLC and mass spectrometry (MS) was performed with a Q Exactive Plus Mass Spectrometer (Thermo Fisher Scientific). Proteins were extracted using the method previously described by Zborowsky & Lindell (10). Briefly, proteins were extracted in 2% sodium deoxycholate and 50 mM ammonium bicarbonate by 2 cycles of sonication, reduced with 3 mM dithiothreitol, modified with 10 mM iodoacetamide, and digested twice with modified trypsin (Promega) at a 1:50 enzyme to protein ratio, in 1% SDC and 50 mM ammonium bicarbonate. Deoxycholate was removed by centrifugation, 1% formic acid was added, and the samples were centrifuged again. The tryptic peptides in the supernatant were desalted using C18 tips (Ultra-Micro), dried, and resuspended in 0.1% formic acid. MS data were analyzed using MaxQuant 1.5.2.8 software against the proteomes of *Synechococcus* WH8109 and the Syn5 phage from the UniProt database. For proteins that were not detected a constant $\log_2$ of LC-MS/MS signal intensity value of 18 was set to replace missing values for statistical analysis.

**Complementation of *E. coli* thioredoxin for T7 infection**

A complementation assay was used to assess whether Syn5 and *Synechococcus* WH8109 thioredoxins complement *E. coli* thioredoxin to enable T7 infection. (11). Serial dilutions of T7 were spotted on *E. coli* expressing different thioredoxins and efficiency of spotting was measured to test complementation as done previously (11). An *E. coli* Δ*trxA* strain lacking the endogenous thioredoxin (Keio collection of single-gene knockout mutants strain ECK3773 (12)) (Table S1) that is necessary for T7 infection was used to express the two non-codon optimized *Synechococcus* WH8109 thioredoxins and the codon optimized Syn5 thioredoxin. Spot assays were conducted to test the efficiency of infection by T7 by pipetting 3 µL of a series of a 10-fold diluted T7 lysate on a lawn of *E. coli* and monitoring for clearings (plaques). Spotting efficiency was compared to *E. coli* Δ*trxA* carrying an empty vector as a negative infection control, and to *E. coli* Δ*lacA* (expressing the endogenous thioredoxin) as a positive infection control.

**Structural comparison of experimentally determined and modeled T7 DNA polymerase and *E. coli* thioredoxin complexes**

Foldseek multimer search (13) was used to find and align modeled complexes of T7 DNA polymerase and *E. coli* thioredoxin to an experimentally determined complex structure (PDB structure accession number 1TKD (14)). Visualization of the alignment was done and imported from the Foldseek web server. The RCSB PDB Pairwise Structure Alignment Tool was used to assess the structural similarity between the complexes by calculating the root mean square deviation (RMSD) and the template modeling score (TM-score) (15). For aligned subunits within the complex, RMSD values range from 0 to infinity and increase with protein size. Lower RMSD values indicate shorter distance variation and thus higher similarities between the subunits (16). The TM-score ranges from 0 to 1 and measures topological similarity with higher values indicating higher confidence that the aligned proteins have the same protein fold. Proteins with TM-score >0.5 are considered to have the same protein fold (17).

### Supplementary Figure Legends

**Figure S1: Multiple alignment of phage and bacterial thioredoxins.** The scale represents the amino acid position from the N-terminus of the gene from *Escherichia coli* strain K12. The putative catalytic site of thioredoxin is surrounded by a dashed red line. Phages are shaded in green and bacteria are shaded in blue. Phage taxonomy and known host are shown in Table S2.

**Figure S2: The effect of Syn5 *trx* deletion on the expression of downstream genes.** (**A**) Relative protein abundance in cells infected by the wild type phage (black) and the *trxA* mutant phage (green). Relative protein abundance values are average $log_2$(LC-MS/MS) intensity values of Syn5 Trx, Cve, and DNA polymerase 1 hour after infection. (**B**) Transcript levels of Syn5 *trx*, *cve* and DNA polymerase genes in wild type and mutant phages. Genes are shown from left to right in their order in the Syn5 genome. All data points are shown, bars show the averages and error bars show the standard deviation of 3 independent experiments. P-values of significantly different MS LFQ intensity or transcript level (paired t-test) are represented by: *p-value<0.05, ***p-value<0.001.

**Figure S3: Comparison of experimentally determined and modeled structures of complexes between DNA polymerase and thioredoxin.** Structural alignment of experimentally determined (yellow) (PDB ID: 1TKD) and modeled structure (blue) of T7 DNA polymerase (red shading) interacting with *E. coli* thioredoxin (blue shading). For T7 DNA polymerase the alignment root mean square deviation (RMSD) is 2.61 angstrom and the template modeling score (TM-score) is

0.94. For *E. coli* thioredoxin alignment RMSD was 0.35 angstrom and TM-score of 0.99. Lower RMSD values indicate a higher structural similarity with values ranging from 0 to infinity and increase with protein size (15). Higher TM-scores indicate higher structural similarity, ranging from 0 to 1 and are less dependent on size compared to RMSD. Proteins with TM-score >0.5 are considered to have the same protein fold (16).

**Figure S4: Optimization of theophylline-dependent riboswitch gene translation in *Synechococcus* WH8109.** Beta-glucuronidase activity after induction with different concentrations of theophylline relative to no theophylline addition (0 mM) at different times after addition: 0.5 h (blue), 7.5 h (orange), 24.5 h (yellow), 48.5 h (purple) and 72.5 h (green). All data points are shown with the bar showing the average and the error bars showing the standard deviation of 4 biological replicates. P-values of significantly different GUS activity relative to 0 mM theophylline over time (repeated measures ANOVA), are shown. * - p-value<0.05, ** - p-value<0.01, *** - p-value<0.001.

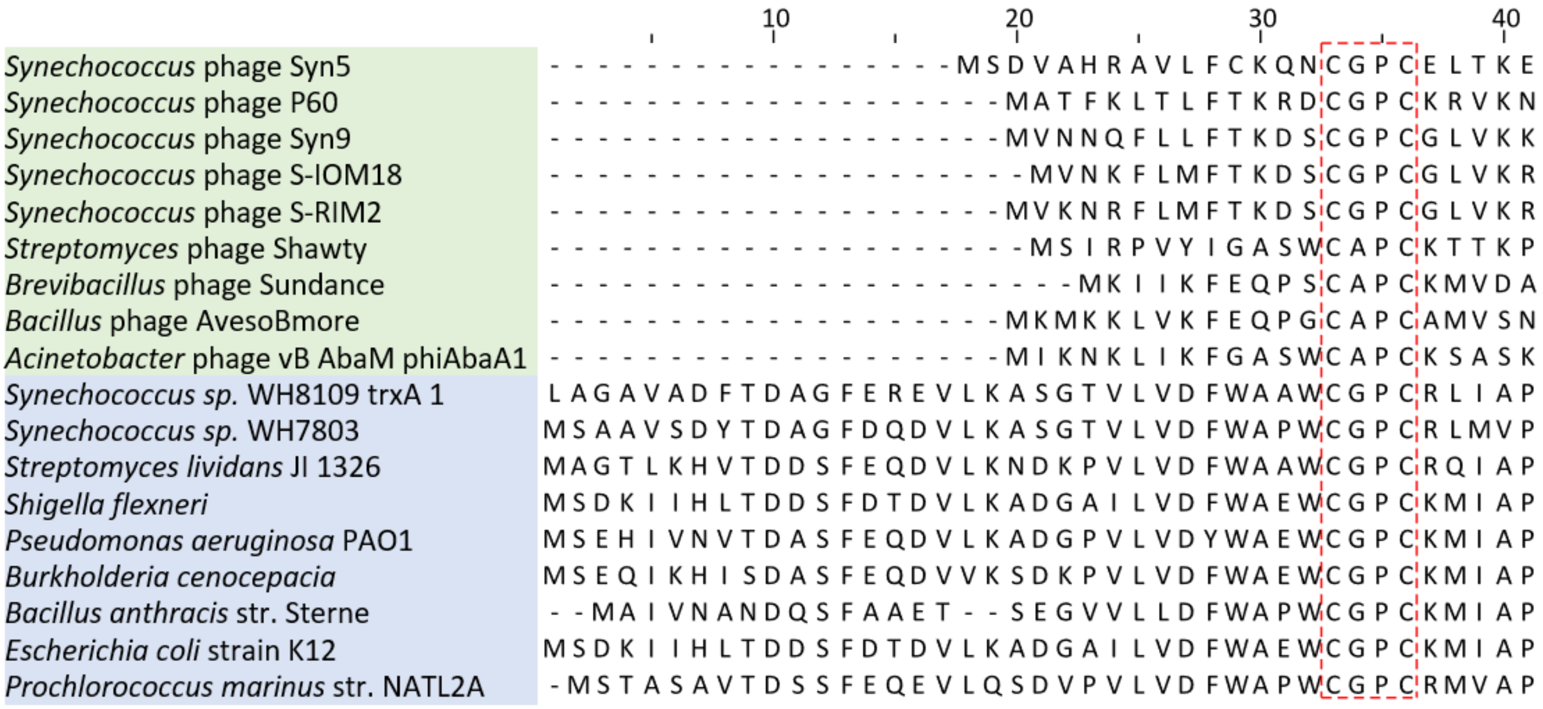

**Figure S1: Multiple alignment of phage and bacterial thioredoxins.** The scale represents the amino acid position from the N-terminus of the gene from *Escherichia coli* strain K12. The putative catalytic site of thioredoxin is surrounded by a dashed red line. Phages are shaded in green and bacteria are shaded in blue. Phage taxonomy and known host are shown in Table S2.

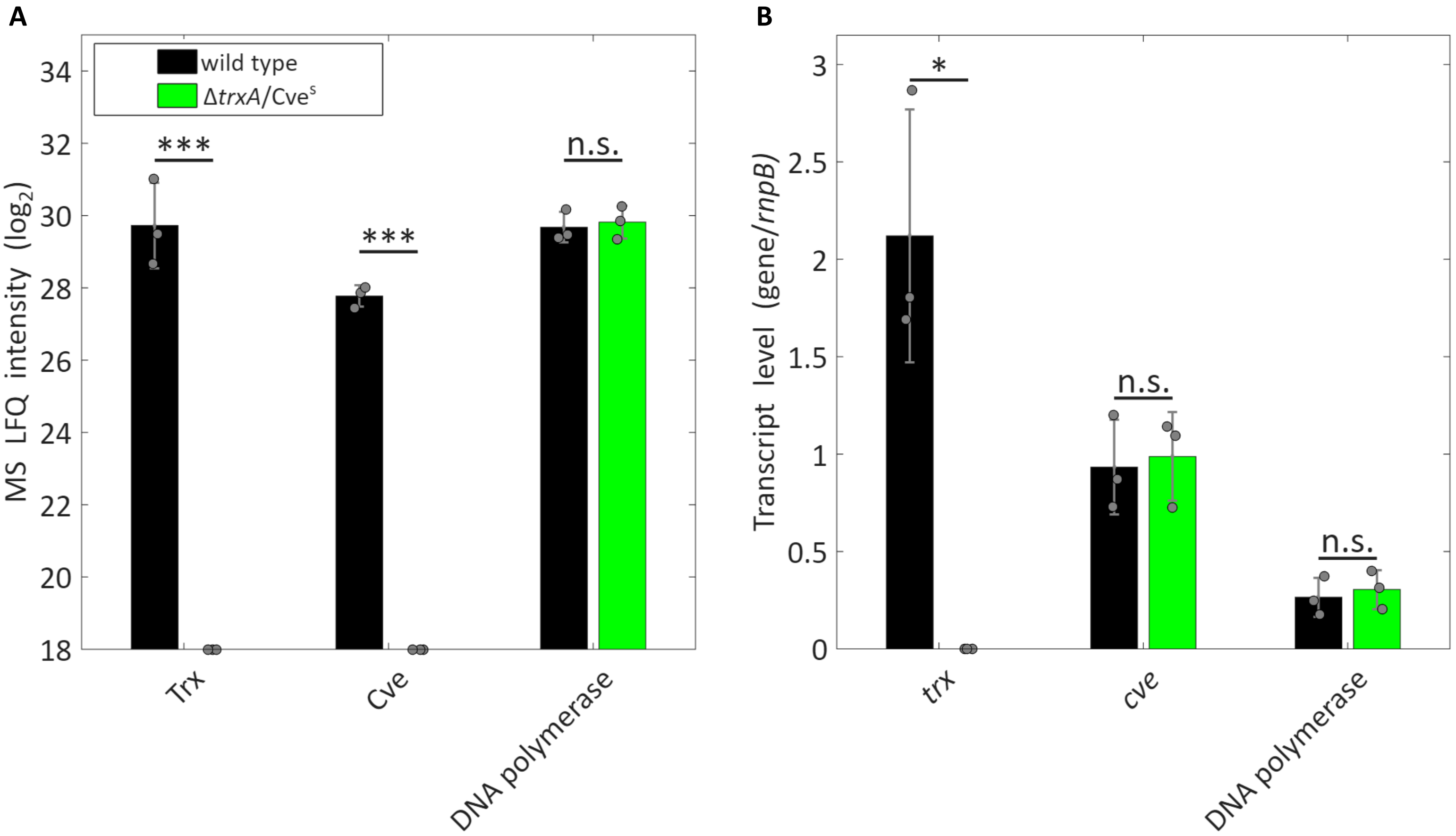

**Figure S2: The effect of Syn5 *trx* deletion on the expression of downstream genes. (A)** Relative protein abundance in cells infected by the wild type phage (black) and the *trxA* mutant phage (green). Relative protein abundance values are average $\log_2$(LC-MS/MS) intensity values of Syn5 Trx, Cve, and DNA polymerase 1 hour after infection. **(B)** Transcript levels of Syn5 *trx*, *cve* and DNA polymerase genes in wild type and mutant phages. Genes are shown from left to right in their order in the Syn5 genome. All data points are shown, bars show the averages and error bars show the standard deviation of 3 independent experiments. P-values of significantly different MS LFQ intensity or transcript level (paired t-test) are represented by: *p-value<0.05, ***p-value<0.001.

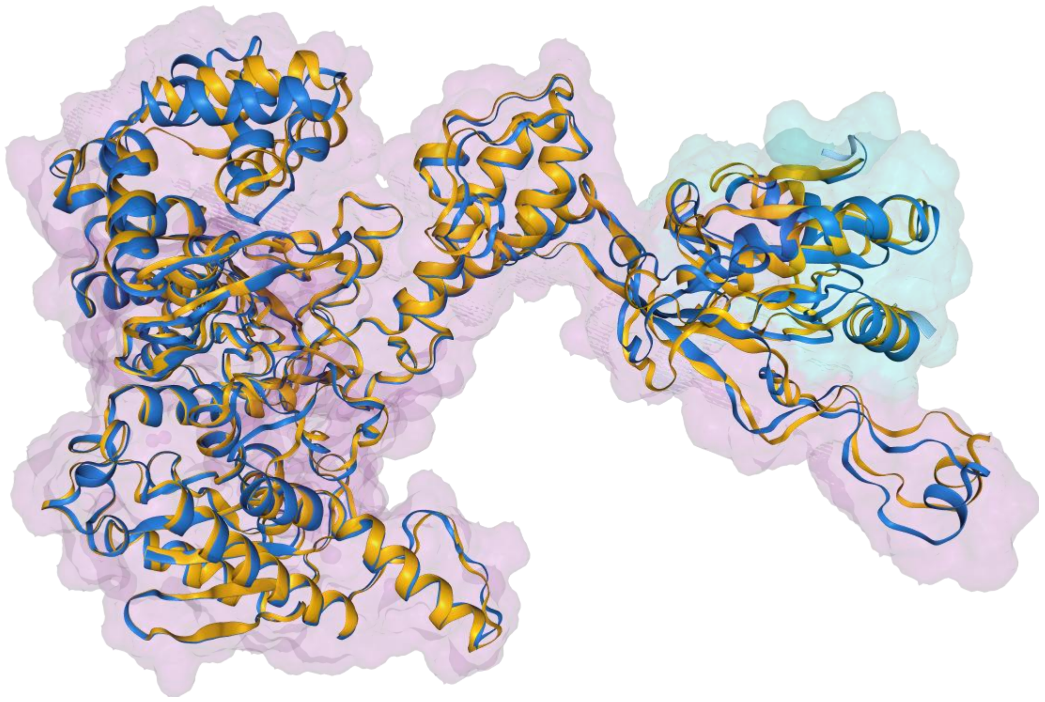

**Figure S3: Comparison of experimentally determined and modeled structures of complexes between DNA polymerase and thioredoxin.** Structural alignment of experimentally determined (yellow) (PDB ID: 1TKD) and modeled structure (blue) of T7 DNA polymerase (red shading) interacting with *E. coli* thioredoxin (blue shading). For T7 DNA polymerase the alignment root mean square deviation (RMSD) is 2.61 angstrom and the template modeling score (TM-score) is 0.94. For *E. coli* thioredoxin alignment RMSD was 0.35 angstrom and TM-score of 0.99. Lower RMSD values indicate a higher structural similarity with values ranging from 0 to infinity and increase with protein size (15). Higher TM-scores indicate higher structural similarity, ranging from 0 to 1 and are less dependent on size compared to RMSD. Proteins with TM-score >0.5 are considered to have the same protein fold (16).

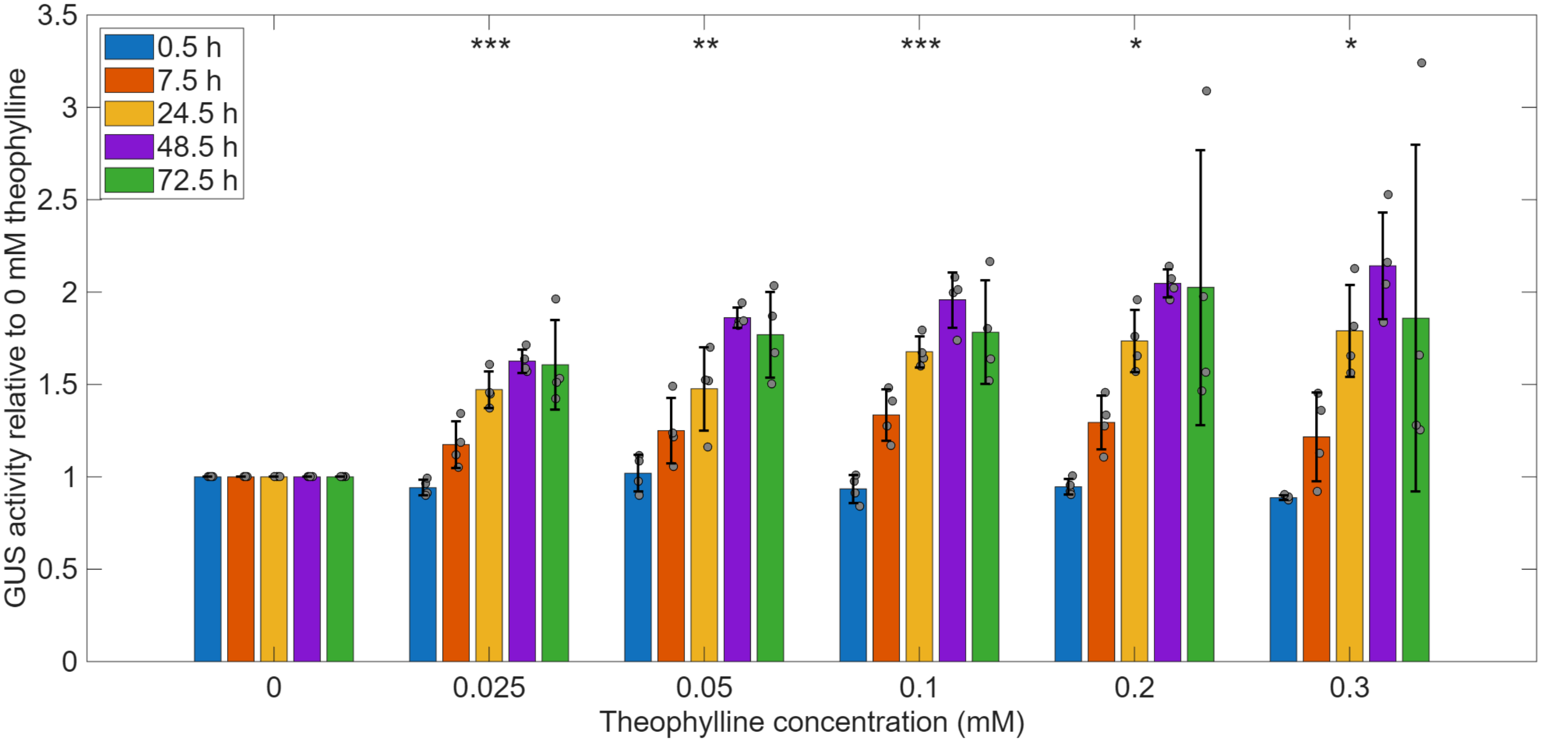

**Figure S4: Optimization of theophylline-dependent riboswitch gene translation in *Synechococcus* WH8109.** Beta-glucuronidase activity after induction with different concentrations of theophylline relative to no theophylline addition (0 mM) at different times after addition: 0.5 h (blue), 7.5 h (orange), 24.5 h (yellow), 48.5 h (purple) and 72.5 h (green). All data points are shown with the bar showing the average and the error bars showing the standard deviation of 4 biological replicates. P-values of significantly different GUS activity relative to 0 mM theophylline over time (repeated measures ANOVA), are shown. * - p-value<0.05, ** - p-value<0.01, *** - p-value<0.001.
